# Nuclear phylogenomics reveals strong geographic patterns in the evolutionary history of *Aloe* and related genera (alooids)

**DOI:** 10.1101/2025.02.28.640761

**Authors:** Yannick Woudstra, Paul Rees, Solofo E. Rakotoarisoa, Ronell R. Klopper, Gideon F. Smith, Nina Rønsted, Olwen M. Grace

## Abstract

**Background and Aims:** With >700 species, *Aloe* and its generic kin (alooids) are a morphologically diverse group of succulent plants with a wide range across Africa, Madagascar and the Arabian Peninsula. Species such as *Aloe vera* and *A. ferox* are cultivated at scale for natural products, whole foods, and cosmetics. Despite substantial α-taxonomy contributions, infrageneric classification of *Aloe* has remained unresolved. Molecular systematics has been compromised by the lack of informative characters in standard markers and high costs of obtaining informative nuclear loci from large genomes (e.g. >15 Gbp); and the difficulty of obtaining quality DNA extractions from material of known provenance. Here these constraints are overcome with target capture sequencing that allows cost-effective sequencing of informative low-copy nuclear loci and unlocks genetic resources from preserved specimens in herbaria as well as silica-dried tissues.

**Methods:** Using a custom kit for alooids, 189 nuclear loci were sequenced in 294 species, including 50 herbarium specimens, to build a new phylogenomic framework for the big genus *Aloe* and 11 closely related genera in the alooid clade. Available taxonomic classifications for *Aloe* sensu stricto were contrasted against the obtained topology in an effort to stabilise taxonomy. Genus-level representations of the family Asphodelaceae were sequenced with the same tool.

**Key results:** The new phylogenomic framework demonstrates the monophyly of the alooids and confirms recent classifications in which smaller genera (*Aloidendron*, *Aloiampelos*, *Aristaloe*, *Gonialoe*, *Kumara*) are separated. Strong geographic patterns in the *Aloe* phylogeny are contrasted by less obvious phylogenetic structure in habit (growth form), and vegetative or reproductive morphology that are mainstays of α-taxonomy.

**Conclusion:** Repeated incidents of adaptive radiation and niche specialisation appear to underlie species diversity in *Aloe*. This study illustrates the power of combined (nuclear) phylogenomic and α-taxonomic inference, including the utility of herbarium genomics, in resolving the systematics of big genera.

**Lay summary:** The alooids are a group of >700 succulent plants, such as *Aloe vera*. Testing their taxonomy with genetics is difficult as *Aloe* genomes are large and very similar between species. A targeted approach was used to classify alooids with highly variable nuclear genes in ±300 species. The evolutionary relationships of alooids are determined principally by geographic distribution patterns, rather than plant morphology or flower shape.

## INTRODUCTION

Molecular phylogenetics has become the framework for integrated systematics (Grace *et al*., 2021). Reconstructed evolutionary history can be contrasted with available taxonomic hypotheses to provide additional evidence for revised classifications. This has led to the presentation of revised systematics for morphologically diverse and taxonomically complex plant families [e.g. Fabaceae (Legume Phylogeny Working Group, 2017), Asteraceae (Mandel *et al*., 2019), Poaceae (Grass Phylogeny Working Group III, 2024), and, at a higher taxonomic rank, flowering plants in general (The Angiosperm Phylogeny Group, 2016)]. In addition, a phylogenetic tree provides meaningful insights about the evolutionary history that gave rise to observed morphological variation (e.g. Zuntini *et al*., 2024). In many plant groups, however, the implementation of molecular systematics has unfortunately been hindered by: (i) a lack of variation in conventional DNA markers, leading to high levels of uncertainty in phylogenetic tree inference (Hollingsworth *et al*., 2011); and (ii) difficulties in obtaining high-quality DNA samples from taxonomically verified reference material. This is particularly the case for highly diverse, “big” genera, i.e. genera containing >500 species (Frodin 2004), which are often characterised by high levels of rapid diversification and comprise many “difficult-to-collect” species, which may require considerable resource and skill to be collected for study.

The emergence of nuclear phylogenomics, particularly with the use of target capture sequencing, has assisted with overcoming these limitations for many plant lineages (Dodsworth *et al*., 2019). With these modern techniques, hundreds of phylogenetically-informative nuclear loci can be sequenced with high coverage at a relatively low cost (Woudstra *et al*., 2022). Target capture sequencing also works well on degraded DNA from historical specimens (Hart *et al*., 2016; Quatela *et al*., 2023), unlocking a “treasure vault” of genetic resources represented by centuries of botanical collecting with global coverage. The inclusion of herbarium specimens, particularly type specimens, is considered the gold standard in molecular systematics (Renner *et al*., 2024) because the application of taxonomic names is based on these specimens, thereby providing the best material with which to test taxonomic hypotheses using molecular data. Target capture sequencing tools are available as universal kits [e.g. Angiosperms353 (Johnson *et al*., 2019)] or clade-specific ranging from family-[e.g. Asteraceae1061 (Mandel *et al*., 2014)] to genus-level [e.g. *Inga* (Nicholls *et al*., 2015)]. Although they are available at a lower cost, universal kits target more conserved loci and may therefore be less suitable for molecular systematics in big genera. In fact, most big-genus phylogenomic studies use a customised approach to target more variable loci [e.g. *Dioscorea* with >600 species (Soto Gomez *et al*., 2019), *Silene* with >1000 species (Quatela *et al*., 2023) and *Begonia* with >2000 species (Michel *et al*., 2022)].

In terms of morphology and ecology, *Aloe* sensu stricto is highly diverse and, in terms of species numbers, qualifies as a “big” genus comprising 591 species (Newton, 2020) with a relatively recent evolutionary history (±15 million years) (Grace *et al*., 2015). *Aloe* and its generic kin (alooids) are a predominantly African group of succulent plants, with a natural geographical distribution range covering much of the continent, as well as Madagascar, the Mascarene Islands, Socotra, and the Arabian Peninsula. Several *Aloe* species are economically important, for example for the medicinal value derived from various fractions of their leaf extracts, supporting a global health products industry, most notably by *Aloe vera* and *A. ferox*. The inner leaf mesophyll tissue (clear gel) of these species is used in cosmetic products, while the leaf exudate of several species, rich in secondary metabolites with antioxidative properties, has been used in traditional medicine to treat, amongst others, indigestion, malaria, and cancer (Grace, 2011; Amir *et al*., 2019). Despite its relatively recent diversification, the genus presents a remarkable level of morphological and ecological variation (Fig. 1), reflected by the publication of names for at least 56 infrageneric groups, some recognised at the ranks of section and series and others informally named morphogroups.

**Fig. 1.**
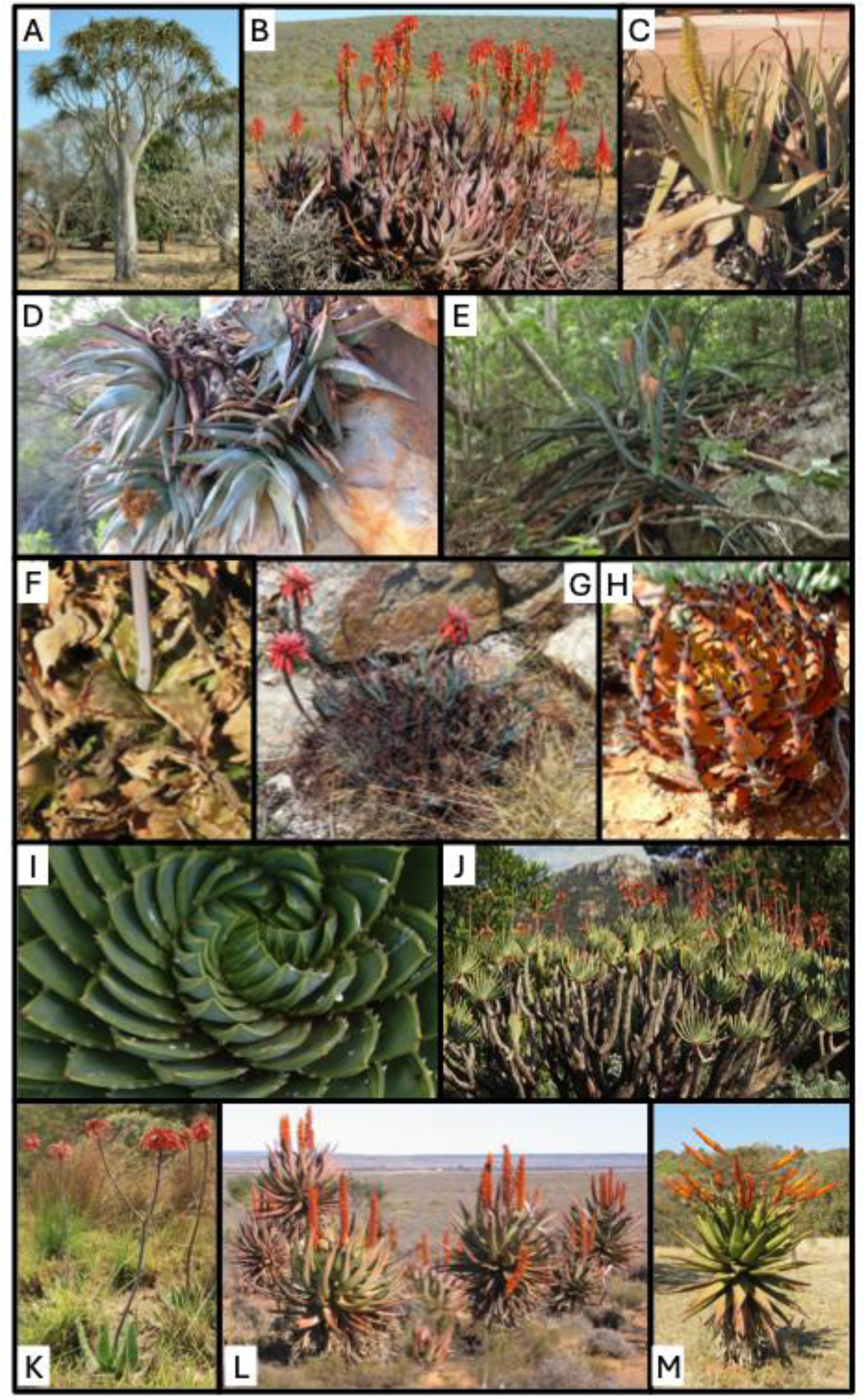
Morphological and ecological diversity of alooid species. Growth form ranges from tall trees (A; *Aloidendron barberae*) to clump-forming shrubs (B; *Aloe framesii*) to small stemless plants (C; *Aloe vera*). Alooids occupy a wide range of habitats (Supplementary Figure S2) ranging from hyper-arid cliff-faces (D; *Aloe meyeri*) to forest understories (E; *Aloe namorokoenesis*). Leaf shape is extremely variable with many species displaying spotted (maculate) patterns (F; *Aloe davyana*), where sometimes the ends of the leaves dry up but stay on. More than 60 species form grass tuft-like leaves (G; *Aloe chortolirioides*) acting as camouflage, whereas others display conspicuous black spines all over the leaves (H; *Aloe melanacantha*). The arrangement of the leaves is usually in a rosette, the most striking example of which is found in the spiral arrangement of *Aloe polyphylla* (I), although distichous (plicate) arrangements are also found (J; *Kumara plicatilis*). Finally, strikingly different inflorescence types are found among the alooids with ball-shaped terminal (capitate) arrangements (K; *Aloe maculata*) and cylindrical racemose panicles (L; *Aloe ferox*) the most common types. Nearly horizontal (oblique) arrangements of inflorescences are also found (M; *Aloe marlothii*). Photo credits: A, I, L – N.R. Crouch; B – A.W. Klopper; C, F, G, J, K, M – G.F. Smith; D – E.J. van Jaarsveld; E – L. Nusbaumer, © Conservatoire & Jardin Botaniques Genève; H – H.M. Steyn.

Although the formal nomenclatural and taxonomic history of *Aloe* dates back to 1753 (Linnaeus, 1753), the first ‘modern’ monograph of the genus was only published more than 150 years later (Berger, 1908). Berger (1908) recognised 181 species that he arranged in more than 30 infrageneric groups; many of these are still upheld today. Berger (1908) based his work on herbarium collections as well as living material cultivated in the Hanbury Garden at La Mortola, Italy. As far as is known, Berger never studied aloes in their natural habitats. Within 20 years following the publication of Berger’s (1908) classification, locally active botanists such as HB Christian (South-tropical Africa), H Perrier de la Bâthie (Madagascar), and GW Reynolds (southern Africa), together described more than 140 species that are still accepted today (Klopper and Smith, 2013). Reynolds produced the most extensive monographs on *Aloe* to date, focusing on southern Africa first (Reynolds, 1950) and later complementing this work with treatments of the *Aloe* taxa of Madagascar and tropical Africa (Reynolds, 1966). In the South African monograph, Reynolds (1950) followed Berger’s classification (1908) but refined and expanded other formally published infrageneric groups. Many of the new species described from tropical Africa and Madagascar, however, did not comfortably fit within the framework of Berger (1908), leading Reynolds (1966) to group them into newly circumscribed morphogroups. In subsequent decades the number of new species discovered and described increased steadily (Klopper and Smith, 2013), with the novel material mainly originating from tropical Africa and Madagascar. By the early 21^st^ century, molecular phylogenetic studies started influencing the classification of the alooids, especially at the rank of genus.

Historically, many genera were segregated from *Aloe*, but these were often eventually reincorporated. Notable alooid segregates reunited with *Aloe* are *Leptaloe* (grass aloes) and *Lomatophyllum* (berried aloes), with *Leptaloe* at present regarded as fitting in *A*. sect. *Leptoaloe*, along with the robust grass aloes, also referred to as slender aloes in the vernacular. The berried aloes are at present treated at the rank of section in the genus *Aloe* (Glen and Hardy, 2000; Grace *et al*., 2013; Newton, 2020). Representatives of *Chortolirion* (bulbous, grass-like aloes), which have a decidedly haworthioid appearance, are now similarly included in an expanded concept of *Aloe* (Manning *et al*., 2014). In contrast, several genera have been reinstated or newly published based on molecular evidence (Grace *et al*., 2013; Manning *et al*., 2014; Grace *et al*., 2015): *Aloestrela* [ancient aloe] (Smith and Molteno, 2019) and *Aristaloe* [awn-leaf aloe] (both monotypic); *Kumara* ([fan aloes] two species); *Aloiampelos* ([rambling aloes] seven species); *Aloidendron* ([tree aloes] six species); and *Gonialoe* ([kanniedood aloes] four species). In addition, the genera *Astroloba* (11 species), *Haworthia* (57 species), *Haworthiopsis* (18 species), *Gasteria* (28 species), and *Tulista* (4 species) are all treated as accepted and distinct from *Aloe* sensu stricto (Newton, 2020). As a result, *Aloe* sensu stricto now includes 591 species (excluding natural and artificial hybrids; CITES Secretariat, in press), many of which have never been placed in any of the infrageneric groups available for the genus.

The taxonomy of *Aloe* and some of its generic kin is therefore in a state of flux. A lack of character variation (and thus resolution) in current molecular phylogenetics (Manning *et al*., 2014; Grace *et al*., 2015; Dee *et al*., 2018), however, resulted in authors of recent taxonomic treatments (Carter *et al*., 2011; Newton, 2020) being reluctant to incorporate at least some of the proposed changes at various generic and infrageneric ranks. Expanding the molecular dataset with plastid phylogenomics (e.g. using whole chloroplast genomes from genome skimming data) can improve this situation, particularly when herbarium material, including type specimens, is incorporated, as evidenced by Malakasi *et al*. (2019) in a study of the tree aloes (*Aloidendron*). For the big genus *Aloe*, however, more variable data from the nuclear genome is needed to track species boundaries (Woudstra *et al*., 2021).

This study aimed to test a highly reduced representation approach to resolve long-standing phylogenetic uncertainties in *Aloe* with new nuclear phylogenomic data from an expanded sampling of species in the genus. A customised target capture sequencing tool (Woudstra *et al*., 2021) was applied to generate an adequate nuclear genomic reference database (Woudstra *et al*., 2024) from which a phylogenomic framework for the alooids (Asphodelaceae subf. Asphodeloideae) was built. The resulting phylogeny was then interpreted taxonomically to inform the stability, and remaining uncertainties, of the classification of *Aloe* and related genera.

## MATERIALS AND METHODS

### Sampling

The basis of this study was formed by sampling preserved (herbarium and living) botanical collections at the Royal Botanic Gardens, Kew (K); Natural History Museum of Denmark, Copenhagen (C); East African Herbarium (EA), the Muséum National d’Histoire Naturelle, Paris (P); and the botanic gardens of the Universities of Potsdam, Uppsala, Göteborg, and Cambridge. The sampling was designed to build (i) a genus-level backbone phylogeny for the family Asphodelaceae and (ii) a comprehensive phylogeny for *Aloe* sensu stricto and related genera (the alooids). Here we apply the term “alooid” to all currently and formerly accepted species and infraspecific taxa of the genus *Aloe* and related genera. These comprise *Aloe* sensu stricto (591 species and 71 additional infraspecific taxa), the smaller genera *Aloiampelos* (seven species and three additional varieties), *Aloidendron* (six species), *Gonialoe* (four species), and *Kumara* (two species), the monotypic genera *Aloestrela* and *Aristaloe*, and the related genera *Astroloba*, *Gasteria*, *Haworthia*, *Haworthiopsis* and *Tulista*. All taxon names referred to in this study follow the checklist provided by World Flora Online (WFO, 2025), unless stated otherwise.

For the Asphodelaceae backbone phylogeny, DNA extracts and ready-to-sequence DNA libraries for one member of each genus in the Asphodelaceae were obtained, except *Kniphofia* and *Haworthia*. These genera were sampled as part of the ongoing Plant and Fungal Tree of Life (PAFTOL) project at the Royal Botanic Gardens, Kew (Baker *et al*., 2022). This included samples from the related alooid genera *Astroloba*, *Gasteria*, *Haworthia*, *Haworthiopsis* and *Tulista*. For *Haworthia*, *H. coarctata* was sampled, which was since placed in *Haworthiopsis* (Rowley 2013) and therefore samples representing *Haworthia* sensu stricto are lacking from this study.

For the other alooids, we used samples from two previous studies (Woudstra *et al*., 2021; 2024) covering *Aloe* (401 samples, 378 species, four additional subspecies and eight additional varieties), *Aloestrela* (one species), *Aloiampelos* (three species), *Aloidendron* (four species), *Aristaloe* (one species), *Gonialoe* (one species), and *Kumara* (one species). The sampling for *Aloe* covered the main taxonomic groups (Berger, 1908; Reynolds, 1950, 1966; Glen and Hardy, 2000), phylogenetic clades (Manning *et al*., 2014; Grace *et al*., 2015; Dee *et al*., 2018), and geographic centres of diversity (Carter *et al*., 2011; Grace et al 2015). Detailed information on sample origins can be found in the Accession Information file (online supporting material). The procedure for sampling followed the methods outlined by Woudstra *et al*. (2024).

### DNA isolation and sequencing

DNA was isolated using either a Qiagen DNEasy kit (for high molecular weight DNA from silica-dried leaf material) using silica columns for purification (Qiagen, Holden, Germany) or using a CTAB protocol (Doyle and Doyle, 1987) combined with a purification protocol optimised for herbarium material (Quatela *et al*., 2023) using AMPure XP beads (Beckman Coulter, Brea, California, USA). For some herbarium samples, a cleaning step was performed before CTAB extraction in an effort to remove high amounts of polysaccharides and polyphenol (Shepherd and McLay, 2011). High-molecular weight DNA samples were fragmented by ultrasonication (Covaris M220, Covaris, Woburn, Massachusetts, USA). Libraries were prepared using NEBNext Ultra II library preparation kits for Illumina sequencing using Multiplex Dual Index sets 1&2 (New England Biolabs, Ipswich, Massachusetts, USA). Pools of 8–27 equimolar indexed library samples were then enriched with the alooid target capture bait panel (Woudstra *et al*., 2021), which uses myBaits^®^ v3 chemistry (Daicel Arbor, Ann Arbor, Michigan, USA).

High-throughput Illumina^®^ paired end sequence data was obtained for the samples described above. As datasets from different studies were combined (Woudstra et al 2021; Woudstra et al 2024, the sequence platform and read length varied from 350 bp PE on Illumina MiSeq to 150 bp PE on Illumina HiSeq. Four samples were generated from Illumina RNAseq data assembled into transcripts (Woudstra *et al*., 2021). Details for each included sample can be found in the sequencing information file (online supporting material).

### Phylogenomics

Sequence data were analysed to produce an updated phylogenomic framework for (i) genus-level infrafamilial relationships for genera in Asphodelaceae and (ii) species-level inter- and infrageneric relationships among species in the alooid clade in the subfamily Asphodeloideae (Fig. 2). For the Asphodelaceae backbone phylogeny, a subsampling with one representative of each genus were used, together with four representatives for the big genus *Aloe*: the southern African *Aloe polyphylla*, the eastern African *A. secundiflora*, the Mascarene *A. tormentorii*, and the iconic *A. vera* that has a strong affinity with Arabian aloes. For the alooid species tree, all suitable samples were used, together with *Bulbine frutescens* as outgroup.

**Fig. 2.**
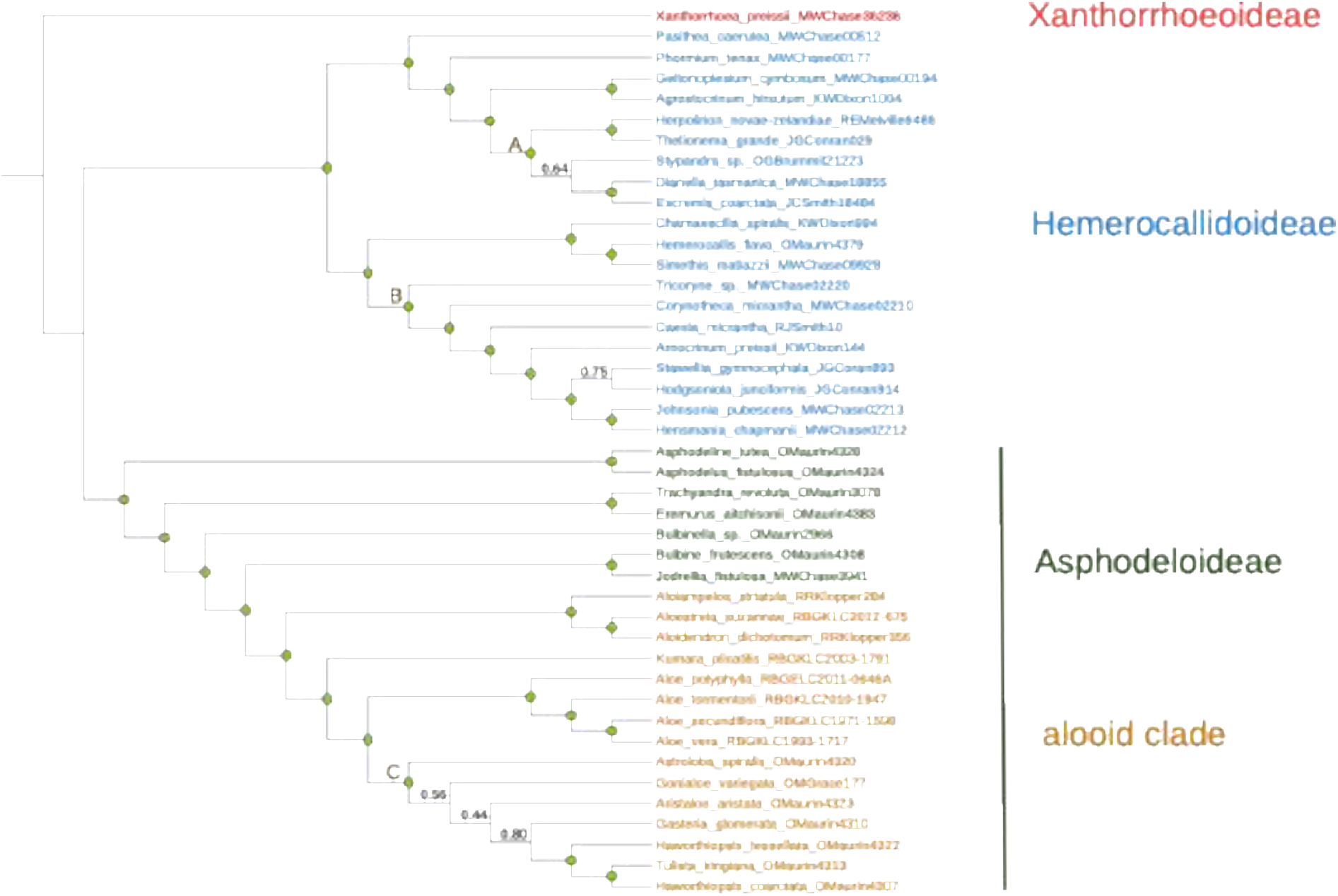
Coalescent summary tree (generated with ASTRAL-III) of the genus-level backbone phylogeny for Asphodelaceae. The tree was manually rooted at the branch for *Xanthorrhoea preissii*, causing the absence of a node support for nodes between subfamilies. Nodes that are fully supported (LPP=1) are indicated with a light green circle and are specified in full if lower (<1). All three defined subfamilies are monophyletic, with the alooid clade being part of the Asphodelaceae subf. Asphodeloideae. The tree can be explored using the following link: https://itol.embl.de/shared/YWoudstra.

A conservative and curated approach was taken to the phylogenomic analysis to minimise issues with sequence alignment and paralogy affecting the inference of phylogenetic relationships. Firstly, information from (overhanging) off-target reads, which can sometimes be assembled in intronic and intergenic regions, were not used as the recovery and coverage of these regions would be relatively random due to the baits being designed to capture exons (Woudstra *et al*., 2021). Secondly, all loci with high rates of paralogy were removed, as identified by the HybPiper paralogy warning script (Johnson *et al*., 2016) and visual inspection of alignments (Woudstra *et al*., 2021). Thirdly, poorly recovered loci (<200 samples) and samples with <50% total recovered target length (≤175 kbp) were excluded. Finally, the resulting phylogeny was curated by removing all samples with questionable phylogenomic placement due to human error in mislabelling of original DNA isolates or in DNA library preparation (Sequencing Information file, online supporting material).

Sequence reads were assembled into contigs and processed into clean, multiple-sequence alignments (MSAs) using the following pipeline: (1) Raw reads were trimmed using Trimmomatic v0.39 (Bolger *et al*., 2014) to remove low-quality (terminal parts of) reads (phred-33 score below 30 - <99.9% accuracy); (2) Trimmed reads were assembled into 189 low-copy nuclear target loci using HybPiper v1.3.1 (Johnson *et al*., 2016) against reference sequences from the original transcriptomes used for the aloe TCS design (Woudstra *et al*., 2021); (3) Assembled target exon sequences were assembled into MSAs using MAFFT v7.429 (Katoh and Standley, 2013); which were subsequently trimmed to remove (4a) insertions (<650 bp) and deletions that are present in less than half of all samples using CIAlign v1.0.14 (Tumescheit *et al*., 2022); (4b) nucleotide sites covered by less than a fifth of all samples using Phyutility v2.2 (Smith and Dunn, 2008); and (4c) previously undiscovered putative paralogous regions by visualisation and manual cleaning in Geneious 9.1 Pro. Sequences <100 bp were removed from the multiple sequence alignments. Woudstra *et al*. (2024) provide more information.

To estimate phylogenetic relationships, maximum likelihood gene trees were generated using IQ-Tree v1.6.12 (Nguyen *et al*., 2015) for each of the loci remaining after the filtering, trimming, and manual cleaning steps. A general time reversible model (GTR) for substitution rates combined with a gamma-distribution for rate heterogeneity (option +G) and a correction for invariable sites (option +I) were deployed. Node support was calculated using 1000 ultra-fast bootstrap replications (option -bb). The 173 resulting gene trees were summarised into a species tree using the coalescent-based model implemented in ASTRAL-III (Zhang *et al*., 2018). This model accounts for incomplete lineage sorting, which is estimated through gene tree discordance, and defined as the ratio of quartets shared between all gene trees and the species tree. The resulting summary trees were visualised and annotated in the interactive Tree of Life programme (Letunic and Bork, 2021). The Asphodelaceae tree was rooted on the branch containing the *Xanthorrhoea preissii* sample, following the Kew Tree of Life (Baker *et al*., 2022) for Asphodelaceae. The alooid species tree was rooted on the branch containing the outgroup sample for *Bulbine frutescens* (subf. Asphodeloideae).

### Taxonomic hypothesis testing

The most comprehensive treatments for *Aloe* were used as taxonomic hypotheses for phylogenetic testing: the monograph of *Aloe* by Berger (1908); the expanded treatments of Reynolds for southern Africa (Reynolds, 1950), tropical Africa (Reynolds, 1966), and Madagascar (Reynolds, 1966); and the floristic treatment of *Aloe* by Glen and Hardy (2000) as part of the *Flora of Southern Africa* series. The groups of Malagasy aloes recognised by Castillon and Castillon (2010) were not tested as the sampling in the present study did not have sufficient coverage of all the groups recognised. Using the taxonomic concepts for sections and series described in these publications, validly published species names were allocated to taxonomic groups (online supporting material). Using the interactive Tree of Life annotation software (Letunic and Bork, 2021) the alooid phylogeny was then annotated with the main infrageneric groups (sections and series) in *Aloe* according to Berger (1908) before adding additional detail from other treatments by Reynolds (1950; 1966) and Glen & Hardy (2000).

Finally, phylogenetic patterns for geographic distribution were tested with eight different areas of diversity distinguished, based on Carter *et al*. (2011) and Grace *et al*. (2015): Southern Africa, including all of South Africa, Lesotho, and Eswatini; Southwestern Africa, including all of Namibia and the dry, southwestern parts of Angola; South Tropical Africa, corresponding roughly to Zambezia including Botswana, Mozambique, Zambia, Zimbabwe, and the eastern parts of Angola; Tropical East Africa, including all of Kenya, Uganda, Tanzania, Rwanda, and Burundi; Tropical West Africa, including the tropical wetter parts of the Congo and the Gold Coast; the Horn of Africa, including all of Ethiopia, Somalia, Eritrea, Djibouti, Sudan, and South Sudan; the Arabian Peninsula, including all of Saudi Arabia, Oman, and Yemen, including Socotra; Madagascar and the Mascarene islands, including La Réunion, Mauritius, Mayotte, the Comoros, and the southern Seychelles.

## RESULTS

This study presents a considerable increase in sequence information for the alooids (Asphodelaceae subf. Asphodeloideae; *Aloe* and related genera) covering all 12 genera (with the exception of *Haworthia* sensu stricto) and 303 taxa (294 species, of which 278 belong to *Aloe*) in total. 142 of these taxa (133 species) have never been sequenced before (Species Information file, online supporting material). The amount of sequence data underlying the phylogeny was considerably increased for all species, from up to seven plastid and nuclear ribosomal markers (previous studies) to 173 low-copy nuclear genes (this study), discounting 16 genes removed due to paralogy or low recovery. The total target dataset (all 189 genes) comprised 350,347 bp compared to 4,693 bp using traditional markers (Woudstra *et al*., 2021).

### A fully resolved backbone phylogeny for Asphodelaceae

The final backbone phylogeny of Asphodelaceae (Fig. 2) comprises all but two (Kniphofia and Haworthia) of the currently accepted genera. Both Hemerocallidoideae (20 genera) and Asphodeloideae (19 genera) are monophyletic, with the monogeneric Xanthorrhoeoideae sister to Asphodeloideae. The alooids (12 genera) comprise a monophyletic clade embedded within subfamily Asphodeloideae. All except five nodes were fully supported (Local Posterior Probability (LPP) >1), with relationships between *Gonialoe*, *Aristaloe*, and *Gasteria* receiving the lowest support.

In Asphodeloideae, *Asphodelus* is monophyletic with *Asphodeline* in a clade that is sister to the rest of the genera. *Trachyandra* and *Eremurus* are also monophyletic, while *Bulbinella* is recovered on a separate branch. *Bulbine* and *Jodrellia* are sister to the monophyletic alooid clade. Within this, *Aloidendron* and *Aloestrela* form a monophyletic clade that, together with *Aloiampelos*, are sister to the rest of the alooid genera. *Kumara* is recovered on a distinct branch, sister to the “core” alooids. *Aloe* sensu stricto is monophyletic and distinct from a clade comprising *Gonialoe* and *Aristaloe* which were previously included in *Aloe* (clade C, Fig. 2). In clade C, *Astroloba* is sister genus to the rest, but the nodes separating *Gonialoe* and *Aristaloe* are poorly supported (LPP=0.56 and 0.44, respectively). *Haworthiopsis tessellata* is separate from *Haworthia coarctata* (i.e. *Haworthiopsis coarctata*), which is monophyletic together with *Tulista*. *Haworthiopsis* is therefore paraphyletic in this topology.

Two monophyletic clades are recovered in Asphodelaceae subf. Hemerocallidoideae. In the smaller clade, the genus *Pasithea* is sister to eight other genera and *Phormium* is sister to the remaining seven. *Geitonoplesium* and *Agrostocrinum* are monophyletic, as are *Herpolirion* and *Thelionema*. The monophyly of *Stypandra* with *Dianella* and *Excremis* is poorly supported (LPP=0.64), while there is full support for monophyly between the latter two. The larger clade involves *Hemerocallis*, which is monophyletic together with *Simethis* and *Chamaescilla*. This group is sister to the former Johnsoniaceae (clade B, Fig. 2), comprising *Tricoryne*, *Corynotheca*, *Caesia*, *Arnocrinum*, *Stawellia*, *Hodgsoniola*, *Johnsonia*, and *Hensmania*. *Hodgsoniola* is monophyletic with *Stawellia* with moderate support (LPP=0.75), while the monophyly of *Johnsonia* and *Hensmania* is fully supported.

### The genera of the alooid clade

The species-level topology for alooids confirms the separation of *Aloestrela*, *Aloidendron*, *Aristaloe*, *Gonialoe*, and *Kumara* from *Aloe* sensu stricto with full support (Fig. 3A). *Aloestrela* is nested within *Aloidendron* and together they form a fully supported monophyletic clade of five species in this topology. *Aloiampelos ciliaris* and *A. striatula* are monophyletic and form a larger monophyletic clade together with *Aloidendron*. *Kumara* is sister to the remaining members of the alooid clade, with the 279 species of *Aloe* sensu stricto forming a large, fully supported monophyletic clade. Although two out of the three sequenced species of *Aloiampelos* are phylogenetically distinct from *Aloe* sensu stricto, *Aloiampelos* is not resolved as a monophyletic clade because *Aloiampelos commixta* is sister to *Aloe* sensu stricto (Fig. 3A).

**Fig. 3.**
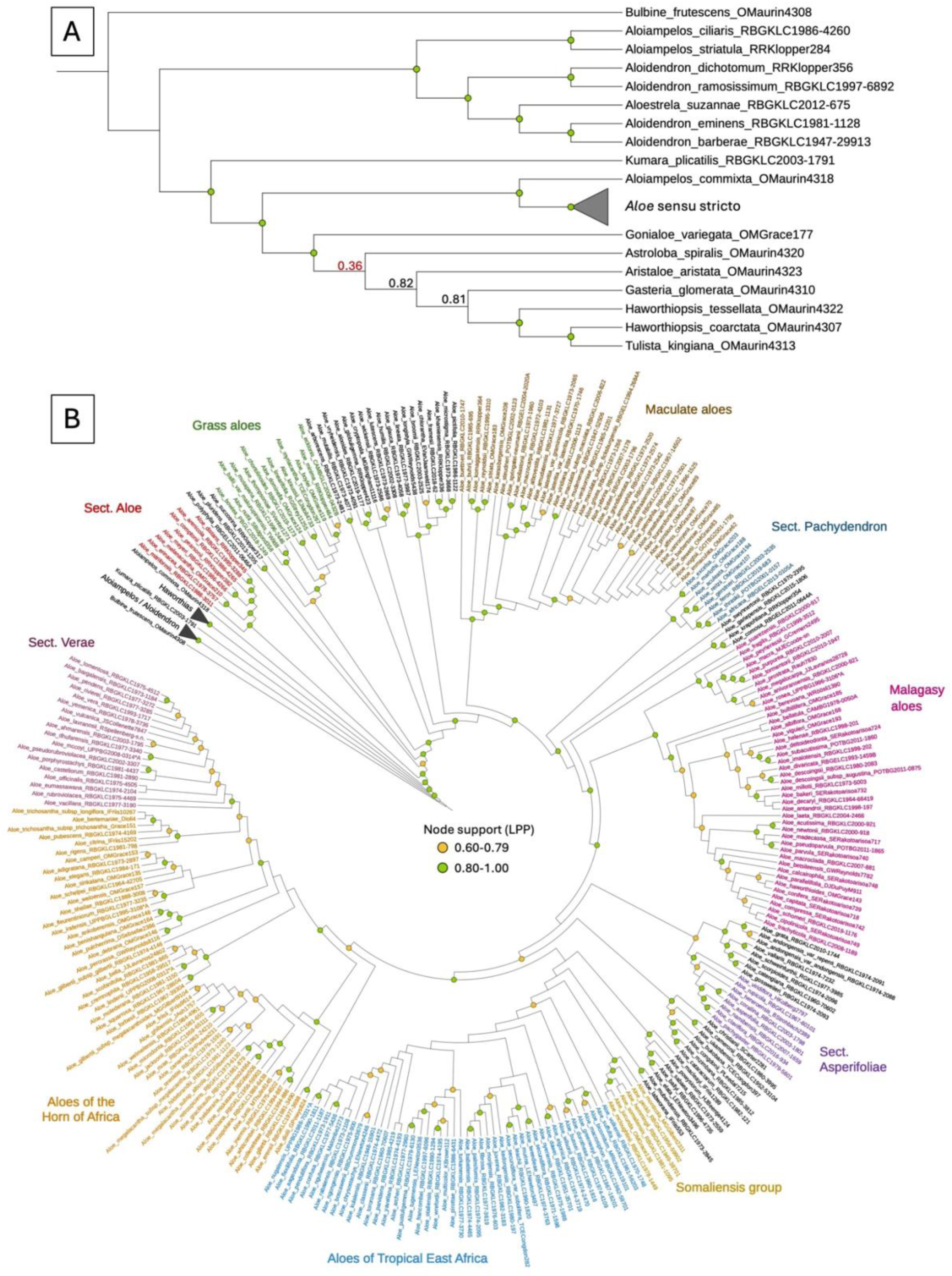
Phylogenetic relationships within the alooids as determined with a coalescent-based summary tree from 173 low-copy nuclear gene trees. (A) Cladogram displaying intergeneric relationships in the alooid clade. (B) Detailed infrageneric cladogram for *Aloe* sensu stricto with labels coloured according to fully supported taxonomic sections (Fig. 4) or geographic distribution (Fig. 5). The tree can be explored using the following link: https://itol.embl.de/shared/YWoudstra.

*Gonialoe* and *Aristaloe* form a fully supported clade with the larger genera *Haworthiopsis* and *Gasteria*, as well as *Astroloba*, and *Tulista*. The genus-level phylogeny of Asphodelaceae (clade C, Fig. 2) and the species-level phylogeny of the alooids (Fig. 3A) disagree about the relationship between *Gonialoe*, *Astroloba*, and the other genera: in the former, *Astroloba* is sister to the rest, while in the latter *Gonialoe* is in this position. In both cases these nodes are not supported (LPP=0.56 and 0.36, respectively) and the sister genus placement therefore remains ambiguous. Both topologies do, however, agree on the placement of *Aristaloe* as sister to *Gasteria*, *Haworthiopsis*, and *Tulista*, with the latter three forming a monophyletic clade.

### Infrageneric relationships in Aloe sensu stricto

The final phylogeny of *Aloe* sensu stricto presented here comprises 279 species with five additional subspecies and three additional varieties (Fig. 3B). Support was generally high (LPP ≥0.80), particularly for the deeper nodes. Gene tree discordance was detected with a normalised quartet score of 0.625, causing low support in some of the shallower nodes of diverse clades (e.g. Maculate aloes). The new topology generally favours the classification of Berger (1908) for the southern African clades (Fig. 4A), whereas the tropical African clades, which are younger, are better characterised by geographic distribution patterns (Fig. 5) than by morphology (Fig. 4B).

**Fig. 4.**
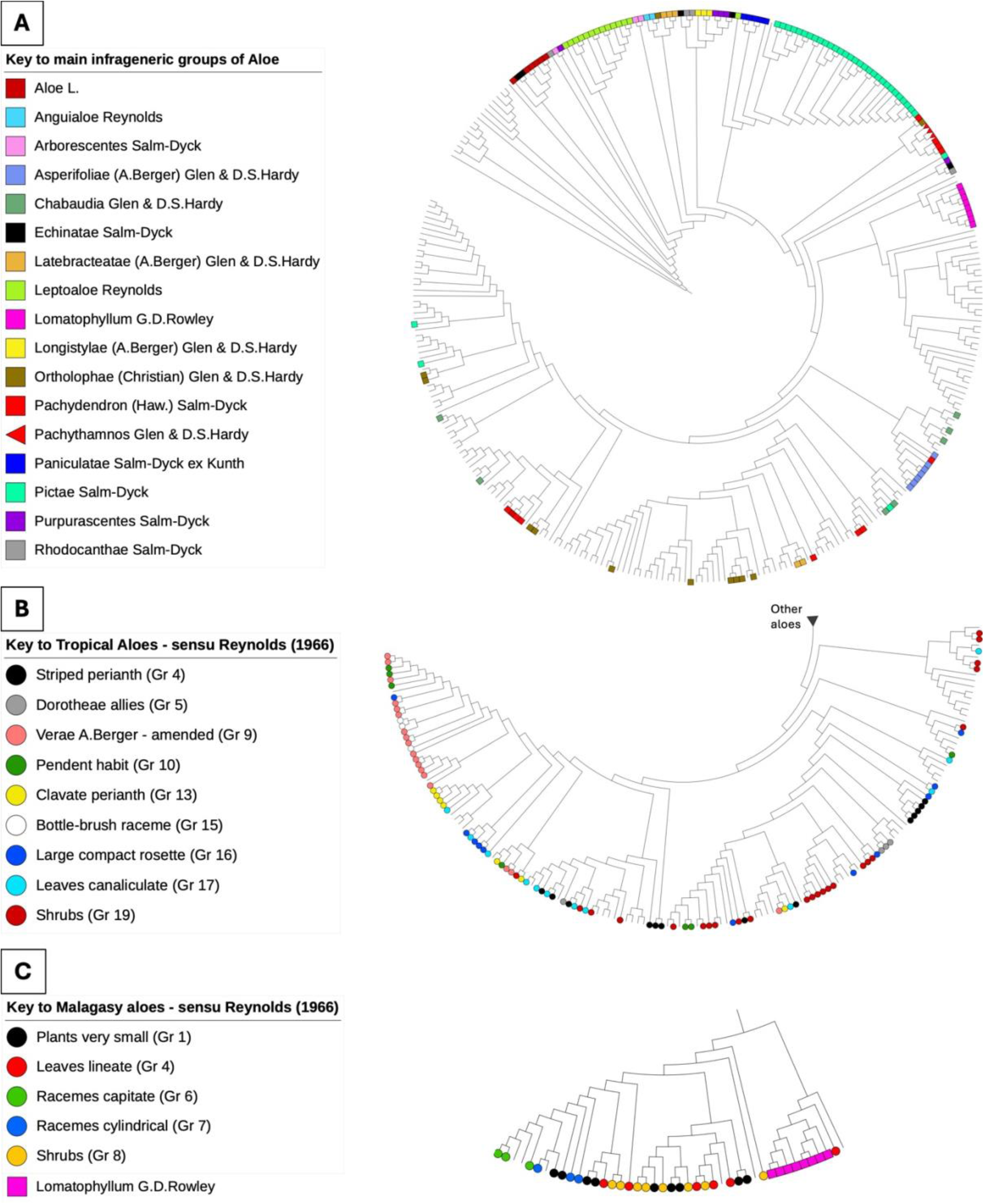
Phylogenetic testing of standing taxonomic hypotheses for the genus *Aloe* sensu stricto. (A) Main sections, following the classification by Berger (1908) with amendments by Reynolds (1950) and Glen and Hardy (2000). (B) Morphogroups for the aloes of tropical Africa and the Arabian Peninsula following the classification by Reynolds (1966). (C) Morphogroups for the aloes of Madagascar and the Mascarene islands, including *Aloe* sect. *Lomatophyllum* as circumscribed by Rowley (1996), and following the classification by Reynolds (1966). Topologies correspond to the coalescent-based summary tree for the alooids (Fig. 3) and can be explored using the following link: https://itol.embl.de/shared/YWoudstra.

**Fig. 5.**
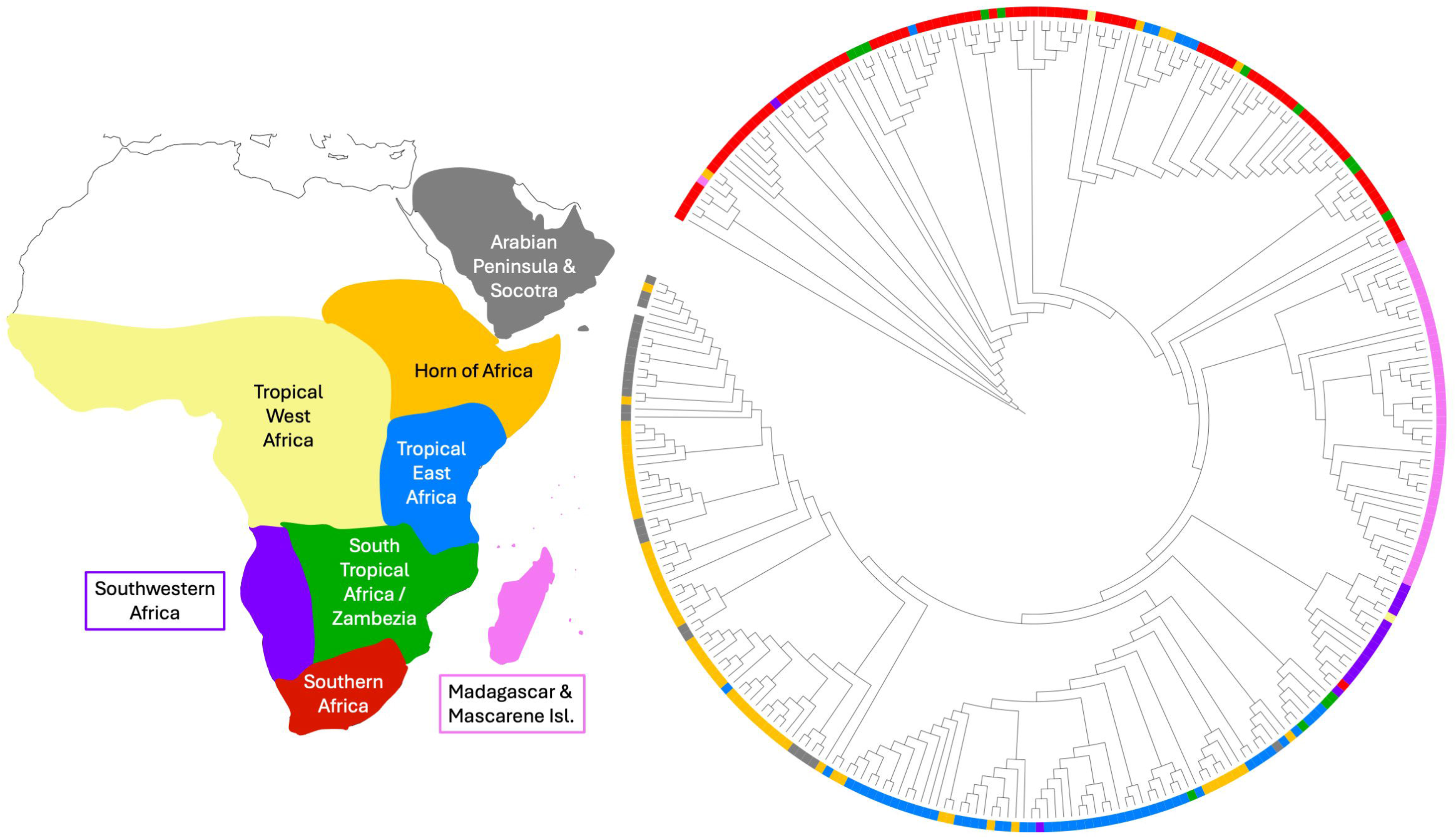
Phylogenomic patterns of geographic distribution in the alooids. Geographic regions are based on the classifications by Carter *et al*. (2011) and Grace *et al*. (2015). Topology corresponds to the coalescent-based summary tree for alooids (Fig. 3) and can be explored using the following link: https://itol.embl.de/shared/YWoudstra.

The older clades are predominantly formed by southern African taxa (Fig. 5). Among these, many of the sections, especially those first included in the monograph of Berger (1908), form clear phylogenetic patterns (Fig. 4A). *Aloe* sect. *Aloe* is sister to the rest of the genus with two species from *A*. sect. *Echinatae* (i.e. *A. erinacea* and *A. melanacantha*) embedded within it. *Aloe* sect. *Leptoaloe* is a monophyletic clade (LPP = 0.73) with the only exception being the bulb-forming *A. buettneri*, which is sister to *A*. sect. *Pictae*. In *A*. sect. *Pictae*, *A*. ser. *Paniculatae* is embedded as a paraphyletic clade. *Aloe* sect. *Latebracteatae* is monophyletic and forms a clade with two paraphyletic members of *A*. sect. *Anguialoe* (i.e. *A. alooides* and *A. vryheidensis*) and *A. globuligemma* (*A*. sect. *Ortholophae*).

With the exception of one species (*A. rupicola*), *A*. sect. *Pachydendron* (single-stemmed, tree-like aloes) is monophyletic and comprised only of southern African species with *A*. sect. *Pachythamnos* embedded within it, along with *A. marlothii* (placed by Berger (1908) in *A*. sect. *Ortholophae*). *Aloe rupicola* (*A*. sect. *Pachydendron*) is instead embedded within the monophyletic *A*. sect. *Asperifoliae*, which comprises species predominantly from Southwestern Africa. Although paraphyletic, the three members of *A*. sect. *Longistylae* (i.e. *A. broomii*, *A. chlorantha*, and *A. longistyla*) are all in the same clade together with three of the five members of *A*. sect. *Purpurascentes* (i.e. *A. framesii*, *A. khamiesensis*, and *A. microstigma*) and two monophyletic members of *A*. sect. *Rhodocanthae* (i.e. *A. glauca* and *A. lineata*). Also included in this clade are *A. humilis* and *A. pictifolia* (from *A.* sect. *Echinatae*), but *A. humilis* resolves as sister to the clade containing the two species of *A*. sect. *Rhodocanthae*, while *A. pictifolia* is sister to *A. microstigma*. *Aloe arborescens* and *A. mutabilis* from *A*. sect. *Arborescentes* are monophyletic, while *A. pluridens* is sister to *A. succotrina* (from *A*. sect. *Purpurascentes*). Another Southern African clade comprises *A. comosa* (from *A.* sect. *Rhodacanthae*), *A. gariepensis* (from *A.* sect. *Purpurascentes*), and *A. krapohliana* (from *A.* sect. *Echinatae*).

While southern African taxa of *A.* sect. *Aloe*, *A*. sect. *Latebracteatae*, *A*. sect. *Leptoaloe*, *A*. sect. *Pachydendron*, and *A*. sect. *Pictae* form strongly supported coherent clades, several sections are polyphyletic: *A.* sect. *Arborescentes*, *A.* sect. *Echinatae*, *A.* sect. *Purpurascentes*, and *A.* sect. *Rhodacanthae*.

The taxonomic sections hitherto recognised in *Aloe* are not recovered intact in the phylogenetic clades of tropical Africa (Fig. 4A). *Aloe* sect. *Chabaudia* (seven species), *A*. sect. *Ortholophae* (10 species), and *A*. sect. *Pachydendron* (six species) are all represented by species spread across three clades covering substantial phylogenetic distances. A similar pattern is found for the three Tropical African members of *A*. sect. *Pictae*, which are spread across two clades. *Aloe* sect. *Latebracteatae* is the only monophyletic section among the tropical African aloes, though separated from the southern African taxa.

All of the tropical African morphogroups recognised by Reynolds (1966) are polyphyletic (Fig. 4B) with only a few groups forming monophyletic clades, but never with all members sampled. Of these, the strongest coherence is found in Group 9 of Reynolds (1966), which is an expanded version of *A*. sect. *Verae*, where only two members are recovered in a separate clade. *Aloe dorotheae* (Reynolds Group 5) forms a monophyletic clade with two of its close relatives but with a third species separate (*A. canarina*). Based on perianth structure, two groups can be formed: those with striped perianths (Reynolds Group 4; clade I, Fig. 4B; five species) that comprise the *A. somaliensis* species complex (Fig. 3B); and those with clavate perianths (Reynolds Group 13; clade II, Fig. 4B; four species).

Geography (Fig. 5) is a better classifier than morphology in *Aloe* sensu stricto, with the Tropical African aloes roughly falling into three geographic clades: Southwestern Africa, Tropical East Africa, and aloes from the Horn of African and Arabian Peninsula. A shrubby growth form is the predominant type of habit among tropical East African species of *Aloe* (24 species, Reynolds Group 19, Fig. 4B; Supplementary Figure S1). The *A. somaliensis* cluster of species mentioned above is nested within the Tropical East African clade and *A*. sect. *Verae* comprises predominantly Arabian aloes. Within *A*. sect. *Pictae*, a monophyletic group is formed by aloes from tropical East Africa and the Horn of Africa.

The Malagasy taxa in *Aloe* sensu stricto form a fully supported monophyletic clade with the monophyletic clade with *A*. sect. *Lomatophyllum* embedded in it (Fig. 4C). As is the case for the tropical African aloes, the Malagasy morphogroups recognised by Reynolds (1966) are not recovered as phylogenetic units. The only coherent group of species from Madagascar comprises three paraphyletic species with capitate inflorescences.

## DISCUSSION AND CONCLUSIONS

### The alooid clade

The findings of this study bring hitherto-elusive phylogenomic evidence to stabilise the current classification of the alooids. The taxonomy of the alooids has been revised significantly in the last three decades, with several genera being split from the core *Aloe* genus (Grace *et al*., 2013; Manning *et al*., 2014; Smith and Molteno, 2019), and older genera being reincorporated (Rowley, 1996; Glen and Hardy, 2000; Grace *et al*., 2013). With a few minor changes, which are detailed below, an intergeneric classification of eleven alooid genera (comprising 730 species) can be stabilised. The present study furthermore provides a framework and genomic tool with which to add previously-uninvestigated species as sequences become available to complete the *Aloe* tree of life.

One of the more easily distinguishable groups of alooids is the tree aloes, i.e. the genus *Aloidendron*, based on their distinctive arborescent, often multi-branched growth form (Malakasi *et al*., 2019). Nevertheless, the systematics surrounding this group has become rather complicated in recent years. Although the arborescent growth form has likely evolved multiple times in the alooids (Supplementary Figure S1), species included in *Aloidendron* are distinct from often single-stemmed trees included in *Aloe* sensu stricto by their height (>3 m), dichotomous branching and non-retention of dead leaves on the stem and branches (Grace *et al*., 2013). For these reasons, the (usually) unbranched Malagasy tree aloe *Aloe suzannae*, which showed phylogenetic affinity with *Aloidendron*, was raised to its own monotypic genus *Aloestrela* (Smith and Molteno, 2019). *Aloe sabaea*, on the other hand, was placed in *Aloidendron* (Manning *et al*., 2014) based on an erroneous record of a 9 m-tall specimen by Reynolds (1966) (Malakasi *et al*., 2019). *Kumara*, the other genus of which one representative attains tree-like dimensions, has distinctly distichous (plicate) leaves. Following the results by Malakasi *et al*. (2019), the present detailed nuclear phylogenomic study of the alooids now reinforces the acceptance of *Aloidendron* and *Kumara* at the rank of genus. *Aloestrela suzannae* is nested well within *Aloidendron* sensu stricto and could be transferred to *Aloidendron*. *Aloe sabaea* is clearly distinct from *Aloidendron* and is related to tropical African tree-like species, such as *Aloe ballyi*, included in *Aloe* sensu stricto, and has already been returned to the genus *Aloe* (Smith *et al*., 2019). *Kumara plicatilis* is also clearly distinct, occurring on a separate branch as sister to the remaining genera in the alooid clade, and the reinstatement of this genus is therefore warranted. Although *Aloidendron pillansii* was not sequenced in this study, its placement in *Aloidendron* was confirmed by Malakasi *et al*. (2019). This species was, however, paraphyletic in that study and thus requires further investigation for its taxonomic status. *Aloidendron tongaense* has yet to be sequenced and investigated phylogenetically and therefore requires further attention to complete systematic studies on *Aloidendron.* At present, the genus *Aloidendron* should therefore comprise seven species, with the inclusion of *Aloestrela suzannae*.

The genus *Aloiampelos* also comprises seven species and is distinguishable by its variously rambling, leaning, or creeping habit, characteristic leaf sheaths and comparatively long internodes (Grace *et al*., 2013). Contrary to *Aloidendron*, this genus is polyphyletic in the present topology (Fig. 3A) with two species showing strong affinity with *Aloidendron* and one, i.e. *Aloiampelos commixta*, being sister to *Aloe* sensu stricto. The placement of this *Aloiampelos commixta* sample was also recovered using the Angiosperms-353 universal target capture kit (Baker *et al*. 2022). In a taxonomic study of *Aloiampelos* by Ellis (2013), *A. commixta* consistently resolved within *Aloiampelos*, often sister to *A. juddii*. However, the various gene trees (based on ITS sequences) presented in that study generally did not show good node support (Ellis, 2013). Further phylogenomic investigation using other accessions, preferably including the type specimen of *A. commixta* and material from *A. juddii*, should confirm whether this taxon does indeed not belong to *Aloiampelos.* Returning this taxon to *Aloe* sensu stricto would be one possible solution, but further investigation is required before any changes to the circumscription of the genus *Aloiampelos* is made.

The kanniedood aloes (genus *Gonialoe*) and the awn-leaf aloe (monotypic genus *Aristaloe*) were separated from *Aloe* based on their phylogenetic affinity with *Haworthia* and *Gasteria* (Manning *et al*., 2014). The results presented here support this separation showing *Gonialoe variegata* and *Aristaloe aristata* as phylogenetically distinct from *Aloe* sensu stricto. *Gonialoe* is based on *A*. sect. *Serrulatae* and now includes four species. To test the monophyly of this small genus of alooids, phylogenomic analysis of the remaining members should be prioritised.

In summary, our data currently support an alooid classification of 11 genera, recognising *Aloe* sensu stricto, *Aloiampelos*, *Aloidendron*, *Aristaloe*, *Astroloba*, *Gasteria*, *Gonialoe*, *Haworthia*, *Haworthiopsis*, *Kumara* and *Tulista* at this rank; with *Aloestrela* incorporated into *Aloidendron*.

### Aloe sensu stricto

The 278 species of *Aloe* sensu stricto included in the present study are monophyletic. The use of nuclear phylogenomics through target capture sequencing has afforded a significant improvement in the molecular systematics of this big genus. Herbarium genomics furthermore allowed the inclusion of many difficult-to-collect and difficult-to-cultivate (e.g. grass aloes) species in the present study, expanding the phylogenetic framework from 197 species of *Aloe* (Grace *et al*., 2015) to 279 out of the 591 currently accepted species. With this expanded phylogenomic framework, it was possible to test existing taxonomic hypotheses based mostly based on vegetative and reproductive morphology using molecular data. The results confirm strong geographic patterns in the evolutionary history of *Aloe*, and the monophyly of five large sections for predominantly southern African taxa and one section for Arabian taxa.

Historically, southern African alooid genera have received more taxonomic attention because of a considerable number of botanists working on the group in the region. Dedicated collecting trips by botanists working with institutes such as the Royal Botanic Gardens, Kew, the South African National Biodiversity Institute, and some South African universities have expanded the available living and preserved collections. As this present study was, for the main part, based on the collections at Kew, good coverage is available for this region. The general emerging pattern is that the infrageneric groupings recognised by Berger (1908), often as they were expanded by Reynolds (1950), are coherent, whereas the groupings published later, e.g. by Reynolds (1966) and Glen and Hardy (2000) are often polyphyletic. Several well-defined groups can be distinguished with, necessarily, their circumscription refined as new taxa are described.

Aptly, the clade that is sister to all other groups recognised in *Aloe* sensu stricto is *A*. sect. *Aloe*, currently comprising the mitre-forming aloes (following the concept of Glen and Hardy, 2000), e.g. *A. arenicola*, *A. mitriformis*, and *A. pearsonii*. These species with their capitate inflorescences are predominantly distributed along the west coast of the Western and Northern Cape provinces of South Africa, as well as in southern Namibia. In the phylogenomic inference presented here, this clade also includes two species, i.e. *A. erinacea* and *A. melanacantha*, both in *A*. sect. *Echinatae*, with sharper, more pronounced teeth on the leaf margins and scattered on one or both leaf surfaces. Like representatives of *A*. sect. *Aloe*, these two species also sucker basally to form clumps of variable density, but their racemes are conical-cylindric and not capitate.

With their grass tuft-like habit and capitate to near-capitate inflorescences, the grass aloes, i.e. *A*. sect. *Leptoaloe*, form one of the most distinctive and visually recognisable clades of aloes. Their association with grasslands and grassy patches on cliffs provides effective camouflage to limit herbivory. These species generally do not succumb to veld fires and are able to resprout from their basally thickened stems and often fusiform roots (Cousins and Witkowski, 2012). These ecological innovations have allowed this group to diversify significantly and it currently includes 62 accepted species (Species and Sections file, online supporting material), >10% of the *Aloe* taxa recognised at present. One grass aloe, *Aloe myriacantha*, has the widest natural geographical distribution of all the aloes, ranging from southern South Africa in a northerly direction to well beyond the equator (Newton, 2020). The grass aloes are clearly monophyletic in the present study with a fully supported node, the only exception being the bulb-forming *A. buettneri*, which resolves here as a member of the “maculate aloes” clade.

The maculate aloes, i.e. *A*. sect. *Pictae*, are the most diverse section in *Aloe*, represented in the present study by 32 out of 90 currently accepted species (Fig. 3B, 4A). Although typical characters of this group, such as (generally) transversely arranged leaf maculations (spots) and flower tubes that are constricted above the ovaries, are not unique to *A*. sect. *Pictae*, they are well defined on a combination of these characters and their leaf surface sculpturing (Grace *et al*., 2009). The nuclear phylogenomic results presented here confirm the monophyly of this group for all southern African species and most of the tropical African species. *Aloe* ser. *Paniculatae*, the coral aloes, is embedded within *A*. sect. *Pictae*. The floral morphology of *A. buettneri* and the coral aloes is similar to those of the maculate aloes, supporting their possible inclusion in *A.* sect. *Pictae*. Although *Aloe swynnertonii* appears on a separate branch in the final phylogeny, the node is not supported (LPP<0.60) and in many of the gene trees this species is part of *A.* sect. *Pictae*, being sister to the other members. The tropical aloes *A. citrina*, *A. congdonii*, and *A. weloensis* are clearly not part of *A*. sect. *Pictae* because they are placed at some distance to the rest of this section (Fig. 4A). Further investigation of the tropical members of *A*. sect. *Pictae* would therefore be warranted to determine their placement as part of the section.

The only section that is monophyletic for both Southern African and tropical African aloes, albeit in two separate clades, respectively, is *A*. sect. *Latebracteatae*, indicating the likely loss and subsequent gain of shared features used to define the section (large floral bracts, flowers on long pedicels (>10 mm), anthers and styles not (or very shortly) exserted from the flower tube; Glen & Hardy 2000). This is a general pattern across the *Aloe* phylogeny, perhaps best exemplified by the multiple separate occurrences of the (generally single-stemmed) tree-habit (>3 m, Supplementary Figure S1). Unbranched trees occur in the Southern African (e.g. *Aloe ferox*), Malagasy (e.g. *A. helenae*), Tropical East African (e.g. *A. volkensii*), and Horn of Africa (e.g. *A. gracilicaulis*) clades. It is therefore not surprising that *A*. sect. *Pachydendron* is polyphyletic.

Other monophyletic groups among the tropical African aloes are *Aloe* sect. *Verae* (Reynolds’ group 9), which includes the popular *Aloe vera*, and a group of small montane aloes from Somalia (Somaliensis group, Fig. 3B). This last group has been treated previously by Carter *et al*. (1984) and includes several members of Reynolds’ group 4 (striped perianth): *A. somaliensis*, *A. hemmingii* (synonymised by Carter *et al*. under *A. somaliensis*), *A. jucunda* and *A. peckii*; with *A. mcloughlinii* and *A. erensii* considered to be close relatives. The nuclear phylogenomic topology in this study clarifies the monophyly of the four species in this group together with *A. mcloughlinii* and the Northern Ethiopian *A. monticola*. The Kenyan *A. erensii*, however, is more distantly related and is monophyletic with *A. diolii* from South Sudan. The monophyly of *A. hemmingii* with *A. somaliensis* would support its synonymy.

### The non-alooid members of Asphodelaceae subf. Asphodeloideae

The arrangement of the other genera in the subfamily Asphodeloideae agrees with previous phylogenetic studies (Seberg *et al*., 2012; Grace *et al*., 2015; McLay and Bayly, 2016) in the monophyly of *Asphodeline-Asphodelus* and the sister relationship of *Bulbine* to the alooid clade. Disagreement is, however, found in the relationships between *Bulbinella*, *Eremurus*, and *Trachyandra*. The present topology agrees with Seberg *et al*. (2012) in placing *Eremurus* and *Trachyandra* together and recovering *Bulbinella* on a separate branch. *Kniphofia* is expected to occupy a separate branch nested among the non-alooid members of the subfamily, according to nuclear data (Baker *et al*. 2022). The monotypic *Jodrellia* is currently synonymised with *Bulbine* (Boatwright and Manning, 2010), a decision that is supported by the data from the present study, although additional species of *Bulbine* need to be investigated to confirm the monophyly of this genus.

### Asphodelaceae subf. Hemerocallidoideae

The present study presents a fully resolved and, with the exception of two nodes, fully supported phylogeny of the monophyletic subfamily Hemerocallidoideae. Several taxonomic and phylogenetic uncertainties are resolved. *Geitonoplesium* is confirmed as a member of this subfamily, sister to *Agrostocrinum*, and *Chamaescilla* is confirmed as sister to a monophyletic clade of *Hemerocallis* and *Simethis* (McLay and Bayly, 2016). The historical polytomy at the base of the clade with *Dianella*, *Excremis*, *Stypandra*, *Thelionema*, and *Herpolirion* (core phormioids, clade A in Fig. 2) is resolved and broadly agrees with the topology presented by McLay and Bayly (2016), with *Thelionema* recovered together with *Herpolirion* and *Dianella* with *Excremis*.

The previously-recognised family Johnsoniaceae forms a monophyletic clade (clade B in Fig. 2). In line with previous phylogenetic efforts (Wurdack and Dorr, 2009; Seberg *et al*., 2012; McLay and Bayly, 2016), *Tricoryne* is sister to the rest of the group in the present phylogeny, followed by *Corynotheca*, *Caesia*, *Arnocrinum*, and the core group formed by *Hodgsoniola*, *Hensmania*, *Johnsonia*, and *Stawellia*. Only one of the aforementioned studies included *Hodgsoniola* (McLay and Bayly, 2016) where it was placed with *Hensmania*. The phylogenomic inference presented here disagrees with this and recovers *Hodgsoniola* with *Johnsonia* instead. *Hensmania* is monophyletic with *Stawellia* with full support. The monotypic *Hodgsoniola* and *Stawellia* (five species) can be united on morphological basis as well, with both genera having basally united tepals (Clifford and Conran, 1998). A possible synapomorphy between *Johnsonia* and *Hensmania* could be the presence of bracts around the flowers.

### Future work

One of the most taxonomically problematic groups in *Aloe*, *A*. sect. *Purpurascentes*, remains polyphyletic in the present phylogenomic treatment. Despite recent work to stabilise the taxonomy of this group (Klopper *et al*., 2023), further adjustment may be necessary. Three of the five representatives of this section included in the present study form a coherent but paraphyletic group: *Aloe framesii*, *A. khamiesensis*, and *A. microstigma*. The phylogenetic distance between the three groupings of species from *A*. sect. *Purpurascentes* supports Klopper *et al*.’s (2023) findings that *A. gariepensis* and especially *A. succotrina* comprise monophyletic clades separate to the rest of the core-Purpurascentes group (i.e. *Aloe framesii*, *A. khamiesensis*, and *A. microstigma*). Based on their morphological analyses, Klopper *et al*. (2023) concluded that *A. pictifolia*, a species regarded by some as a relative of *A. microstigma*, probably does not belong to the *A*. sect. *Purpurascentes*, despite some morphological similarities that might support such a relationship. However, *A. pictifolia* was not included in the molecular analysis of Klopper *et al*. (2023) and in results from the present study, it resolves as sister to *A. microstigma*. Its sectional placement could therefore be reconsidered. A detailed study of all ten species and infraspecific taxa currently included in this section, as well as *A. pictifolia*, using our target capture tool may suggest a refined classification for *A*. sect. *Purpurascentes*.

Another predominantly Southern African section that requires further investigation is *A.* sect *Echinatae*, which clearly does not consist of a natural grouping of species based on molecular data. Members of this section resolve in four different phylogenetically distant clades in the phylogenomic results presented here. As mentioned above, the placement of *A. erinacea* and *A. melanacantha* within the clade comprising *A*. sect. *Aloe*, and the sister-relationship of *A. pictifolia* and *A. microstigma* (from *A*. sect. *Purpurascentes*) could be morphologically justified. More difficult to explain from a morphological perspective is the placement of *A. humilis* as sister to *A. glauca* and *A. lineata* (from *A.* sect. *Rhodacanthae*), as well as the placement of *A. krapohliana* in a clade with *A. comosa* (from *A*. sect. *Rhodacanthae*) and *A. gariepensis* (from *A*. sect. *Purpurascentes*).

Beyond Southern Africa, there are three coherent monophyletic groups: *A*. sect. *Asperifoliae*, comprising arid-adapted aloes distributed in Southwestern Africa; the *A. somaliensis* species complex, comprising small aloes from the Horn of Africa; and *A*. sect. *Verae* comprising mainly Arabian aloes, including the iconic *Aloe vera*. The origins of *A. vera* have long been enigmatic (Grace *et al*. 2015) as it is only known from cultivation and individuals that have likely escaped from cultivation. The prolific plantations and wild growth of *A. vera* in the Caribbean have led to the popularisation of the now synonymous name *A. barbadensis* (Grace 2011). Taxonomically, however, it has often been recognised as a close relative of extant wild species on the Arabian Peninsula and the placement of *A. vera* in *A*. sect. *Verae* along with other Arabian aloes in the present study supports this hypothesis.

To further investigate the *Dianella*, *Excremis*, *Stypandra*, *Thelionema*, and *Herpolirion* clade in Asphodelaceae subf. Hemerocallidoideae, it would be useful to sequence the monotypic genus *Rhuacophila*, which is often synonymised with *Dianella* but shows strong affinity with *Stypandra* (Wurdack and Dorr, 2009; McLay and Bayly, 2016).

### Conclusion

The customised target capture sequencing method has proved highly suitable as a pragmatic solution to expand sequencing efforts in the alooids, despite the large numbers of species and large genome sizes. Strong geographic patterns in the present evidence provide a key to further resolve the infrageneric classification of *Aloe*. Reclassification of *Aloe* sensu stricto will therefore have to build on the approach of Reynolds (1950, 1966): with separate treatments for Southern Africa, Madagascar, and tropical Africa. Much work is still needed for Madagascar and tropical Africa, where the existing morphogroup classifications cannot be upheld (Fig. 4B & 4C) and where many recently described species have yet to be sampled and sequenced. A collaborative approach is therefore advocated, involving local botanists and large regional herbaria, such as those of the South African National Biodiversity Institute (SANBI) in Pretoria (PRE) and Cape Town (NBG) for southern Africa and Madagascar, the East African Herbarium (EA) for tropical East Africa and Tsimbazaza Herbarium (TAN) for Madagascar, to achieve a comprehensive revision to the infrageneric classification of *Aloe*.

## AUTHOR CONTRIBUTIONS

Conceptualisation: O.M.G., Y.W., & N.R.; Specimen curation: P.R., O.M.G., R.R.K., G.F.S. & S.E.R.; Data generation (DNA sequences): Y.W.; Data curation (taxonomic): R.R.K., G.F.S.; Data curation (traits): Y.W.; Analysis (phylogenomic inference): Y.W.; Manuscript writing: Y.W. with help of all co-authors.

## SUPPLEMENTARY DATA

Supplementary Data is available in the PDF file named “Supporting Information” and contains two graphs displaying alooid topologies annotated with habit (growth form) and habitat, respectively.

## ACKNOWLEDGEMENTS

The authors thank collectors, horticulturalists and herbarium curators at the different botanical institutes that provided provenanced reference material to the study: Royal Botanic Gardens, Kew, and Kew Herbarium (K); Royal Botanic Garden Edinburgh (E); East Africa Herbarium (EA); Herbarium of the Musée National d’Histoire Naturelle, Paris (P); University of Uppsala Herbarium (UPS) and Uppsala Botanic Garden, Gothenburg Botanic Garden; Potsdam Botanic Garden; and Cambridge University Botanic Garden. We thank Anne-Sophie Quatela for advice on DNA extractions from herbarium material; and Kennedy Matheka for providing material of *Aloe ngutwaensis* before the official publication of the species description. Computational resources were provided by UNINETT Sigma2 - the National Infrastructure for High Performance Computing and Data Storage in Norway.

## DATA AVAILABILITY STATEMENT

Raw sequencing data used in this study are deposited in the Sequence Read Archive (SRA) of the U.S. National Center for Biotechnology Information (NCBI) under Bioprojects PRJNA1120847 (for alooids), PRJNA1122593 (for other members of the Asphodelaceae), and PRJNA1120785 (for data from the alooid target capture tool design study (Woudstra *et al*., 2021)). Other data related to this manuscript are available through online supporting material deposited in FigShare at DOI 10.6084/m9.figshare.28435394. These include (i) the reference sequences for the four species used in the design of the alooid target capture tool, (ii) Accession Information file, (iii) Sequence Information file, (iv) Species Information file containing info on the taxonomy, geographic distribution and growth form of species investigated in this study, (v) Species-and-Sections file containing taxonomic information on all *Aloe* names published to date, and (vi) a directory containing the assembled sequences, cleaned alignments, gene trees, species trees and the code to produce these.

