## Supporting Information for "Nuclear phylogenomics reveals strong geographic patterns in the evolutionary history of *Aloe* and related genera (alooids)"

**Supporting information to Woudstra et al., 2024: "Nuclear phylogenomics reveals strong geographic patterns in the evolutionary history of alooids (Asphodelaceae subfam. Alooideae)"**

Contents:

Fig. S1 – Phylogenomic patterns in life form across Asphodelaceae subfam. Alooideae
Fig. S2 – Phylogenomic patterns in habitat across Asphodelaceae subfam. Alooideae

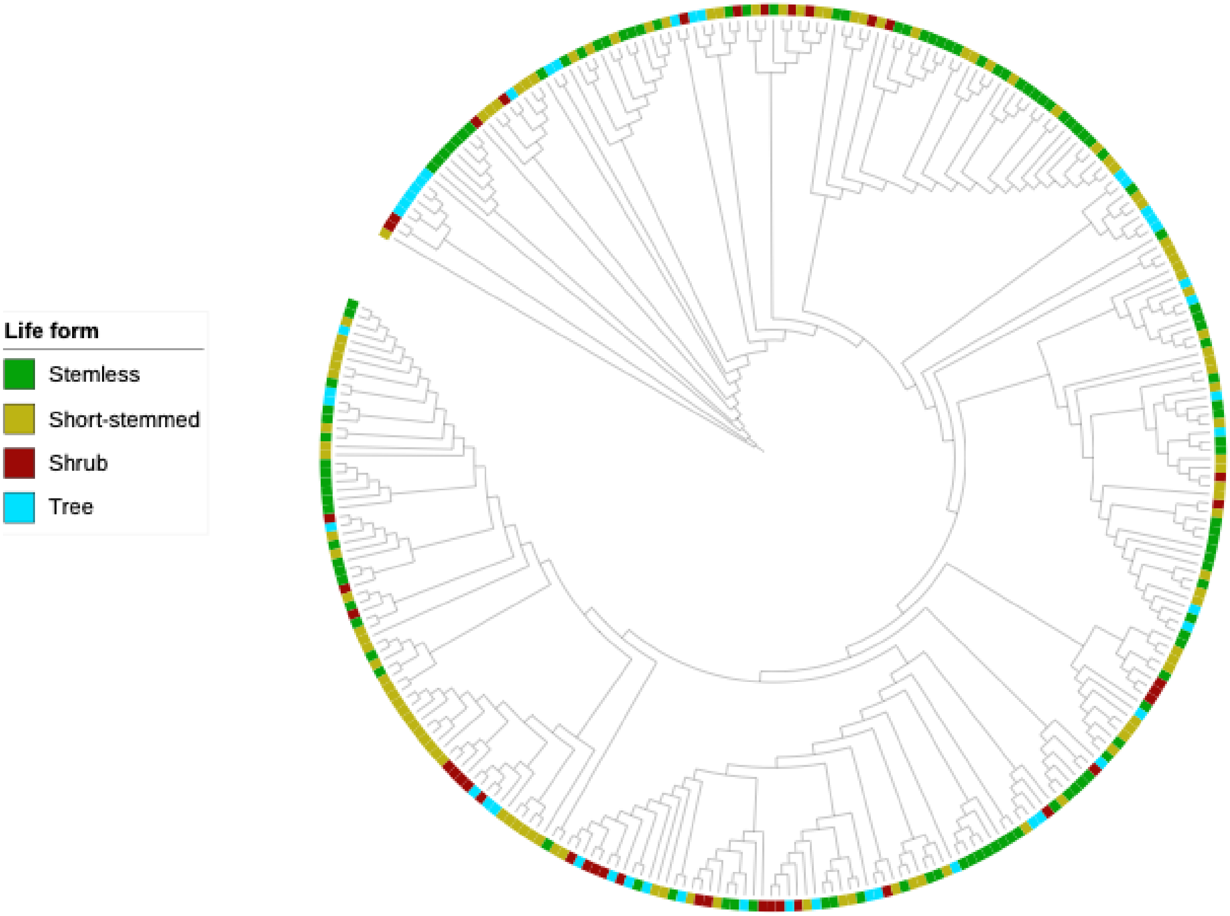


**Fig. S1** Phylogenomic patterns in life form across Asphodelaceae subfam. Alooideae. Trait data is based on species descriptions listed in the overview by Newton (2020). Different life forms are defined as follows: Stemless – plants without stems; Short-stemmed – plants with singular unbranched stems up to 150 cm above the ground; Shrub – plants with branched stems up to 200 cm above the ground; Tree – plants with erect stems (branched or unbranched) at least 200 cm above the ground. Tree topology is based on the ASTRAL-III summary species tree for Asphodelaceae subfam. Alooideae (Fig. 3B of the main manuscript), which can be explored using the following link: https://itol.embl.de/shared/YWoudstra.

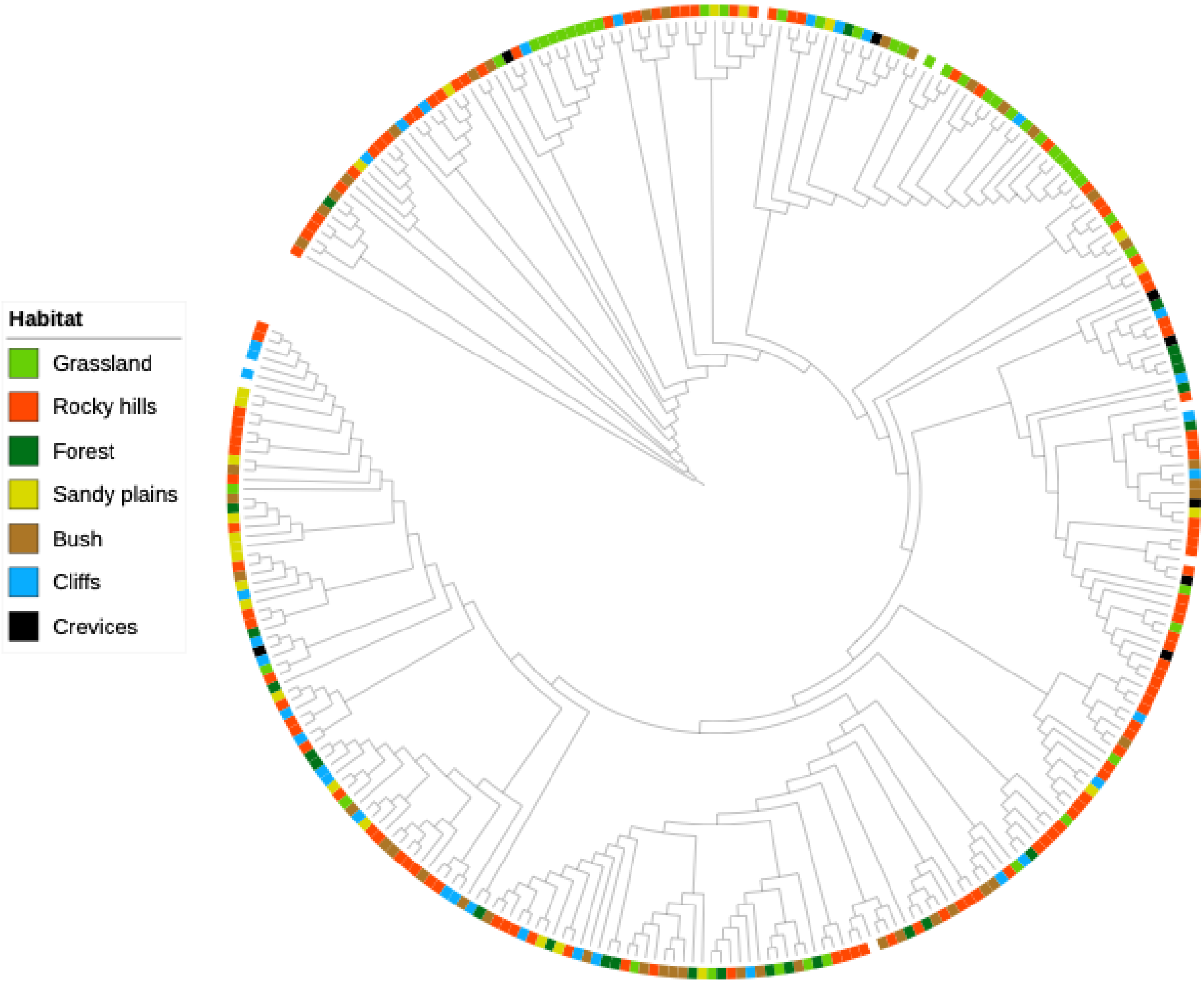


**Fig. S2** Phylogenomic patterns in habitat across Asphodelaceae subfam. Alooideae. Habitat data is based on species descriptions listed in the overview by Newton (2020). Tree topology is based on the ASTRAL-III summary species tree for Asphodelaceae subfam. Alooideae (Fig. 3B of the main manuscript), which can be explored using the following link: https://itol.embl.de/shared/YWoudstra.
